# Tendon Inversion Improves Tendon-to-Bone Healing in a Rat Bicep Tenodesis Model

**DOI:** 10.1101/2024.05.31.596842

**Authors:** Ting Cong, Thomas M. Li, Dustin C. Buller, Varun Arvind, Philip Nasser, Damien M. Laudier, Harrison R. Ferlauto, Arielle J. Hall, Deborah M. Li, Alice H. Huang, Paul J. Cagle, Leesa M. Galatz, Michael R. Hausman

## Abstract

**Purpose:** Tendon- or ligament-to-bone repair remains a surgical challenge. While bone tunnel fixation is a common surgical technique whereby soft tissue is expected to heal against a bone tunnel interface, contemporary methods have yet to recapitulate biomechanical similarity to the native enthesis. In this study, we aim to demonstrate that inside-out longitudinal tendon inversion may improve bone tunnel healing with the hypothesis that inversion removes the gliding epitenon surface from the healing interface thereby improving tunnel interface healing.

**Methods:** 40 male Sprague-Dawley rats underwent either native tendon tenodesis (control group) or tendon inversion tenodesis (experimental group). Interface tissue was harvested 8 weeks post-operatively. Biomechanical testing was performed to assess tensile strength and modes of failure. Histology was performed to assess tissue architecture, and immunohistochemistry was used confirmed abrogation of epitendinous lubricin from interface tissue.

**Results:** Neither surgical intervention led to discernible adverse effects on animal health. Maximum tensile strength increased after tendon inversion compared to control surgery. The extracellular matrix protein lubricin was reduced with tendon inversion, and specimens with tendon inversion had greater healing scores and collagen fibril alignment at the healing interface.

**Conclusions:** Tendon inversion improves bone tunnel healing in rats.

**Clinical Relevance:** Our findings suggest that longitudinal tendon inversion, or inverse tubularization, in a rat biceps tenodesis model improves tendon-to-bone healing in part due to disruption of the epitendinous surface at the bone healing interface. This work provides molecular insight into future improvements for tendon-to-bone repair surgical techniques.

## INTRODUCTION

Tendon-to-bone repair (surgical tenodesis) is central to many orthopaedic procedures, including anterior cruciate ligament reconstruction (ACLR), rotator cuff repair (RCR), flexor and extensor tendon avulsion injuries, and tendon transfers of the upper and lower extremities. The hard-to-soft tissue transition formed during repair largely consists of disorganized scar, a healing adaptation by which the body distributes load upon a larger surface area^1^. To date, there is no surgery that recapitulates the native strength of the tendon-bone interface. Though this may explain re-rupture rates seen in many orthopaedic soft tissue procedures^2^, no orthopaedic practitioner can predict which patient is at risk for re-rupture, and there has yet to be a method to regenerate the functional native transitional tissues of the enthesis^3^. As such, there is a need to further study mechanisms of tendon-to-bone healing.

Current techniques for tendon-to-bone repair follow surgical dogmas rather than biological ones. Specifically, if a tendon has redundant length, repair via a bone tunnel or socket is possible as this approach maximizes surface area by which a tendon contacts the bone. If a tendon does not have redundant length, as seen in RCR, it is repaired by surface tenodesis using suture anchors or transosseous sutures^3^. Literature suggests that mechanical strength of the repair may largely be aperture-based rather than length-based^4^. This implies that biological healing for tendon-to-bone repair occurs primarily at the periosteal juncture which, with recent evidence suggesting that the periosteum harbors a special population of skeletal stem cells^5^, points to the importance of soft tissue contribution in tendon-to-bone repair.

While previous studies have focused on surgical technique or implant design, less has been asked about the tendon tissue itself. The majority of flexor tendons, proximal biceps tendons, and hamstring tendon grafts are micro-anatomically similar. Tendons are designed to slide and consist of an epitendinous layer expressing extracellular lubricin (proteoglycan 4; PRG4), among other molecules, to assist in tendon lubrication^6^. Further, tendons act as a bundle of tendon fascicles that slide against themselves^6^. By extension, surgical reliance on epitendinous healing against bone tunnel walls may be limited by 1) structural and protein expression features of the gliding epitenon at the healing interface and 2) disengagement of endotendinous fascicles in traditional bone tunnel healing.

Here, we test the hypothesis that inside-out tendon inversion improves tendon-to-bone healing in a rat long head of the biceps (LHB) tendon-to-bone tunnel repair model. We find that this procedure does not produce adverse effects and results in a biomechanically stronger tenodesis. We also find that tendon inversion improves bone tunnel healing by histologic scoring, decreases interface lubricin expression, and increases interface collagen fibril alignment. These data provide insight into soft tissue manipulation in surgical tenodesis and suggest novel considerations in tendon-to-bone repair.

## METHODS

### Study design

To address our clinical question, we used a LHB tendon (“biceps tendon”) tenodesis model. We chose this system in rat as tissue size and length are amenable to surgical manipulation without involving larger research animals. The surgical protocol mimics biceps tenodesis, a procedure commonly performed in humans where bone tunnel fixation is commonly used. Further, rat biceps anatomy is similar to that of humans, allowing for translational potential.

### Animals

40 male Sprague-Dawley rats of 12 weeks of age were used. All rats were purchased from Charles River Laboratories. Rats were kept in a specific pathogen-free barrier facility, and all animal procedures including protocols to reduce pain, suffering, and distress were performed in accordance with the regulations of the Institutional Animal Care and Use Committee (#AR202300000107). 20 rats underwent unilateral biceps tenodesis without tendon inversion (control group), and 20 rats underwent unilateral biceps tenodesis with tendon inversion (experimental group). 14 animals from each group were dedicated to biomechanics, and 6 were used for histologic analyses. Animal numbers were calculated based on a power analysis using an effect size estimation based on prior studies^7^.

### Surgical method

After isoflurane induction, rats were placed in a supine position. The forequarter was shaved, sterilely prepared, and draped. A 2 cm incision was made anteriorly along the long axis of the humerus, centered at the level of the humeral head. Subcutaneous tissue was spread until the trapezius, clavicle, deltoid, and pectoralis were visible in the surgical field with exposure carried distally to the level of the deltoid tuberosity.

Next, the deltopectoral interval was released (Figure 1A). Iris scissors were used to open the interval by spreading. The anterior deltoid origin was cut along its clavicular origin with care taken to leave a small cuff of tissue on the clavicle to facilitate later repair.

**Figure 1.**
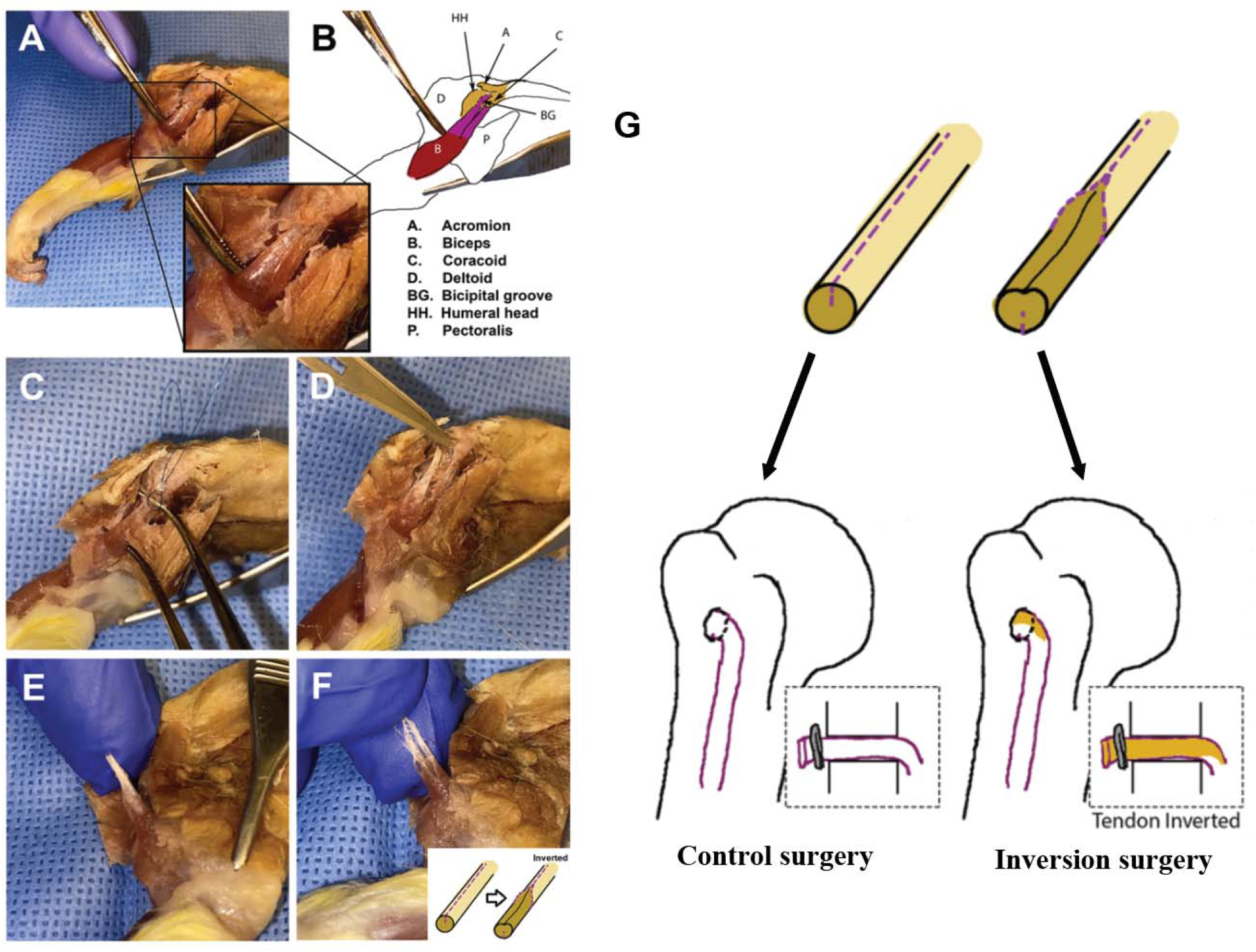
Surgical approach to the rat biceps tendon. (A) A deltopectoral approach with anterior deltoid release was used to expose the biceps (inset). (B) A schematic drawing depicts the anatomic structures. (C) The LHB is tagged and released from its glenoid attachment at the upper extent of the transhumeral ligament. (D) The LHB can alternatively be sharply released from the glenoid. (E-F) The tendon can be split lengthwise to invert the tendon inside-out (“Inverse tubularization”, inset). (G) Summary schematic.

The LHB was not immediately apparent at this time, though the long head musculotendinous junction can occasionally be seen through the overlying groove tissue. The upper falciform ligament was released, and sharp dissection was continued while staying centered on the humerus until the distal tendon of the LHB was be visualized. The groove was then opened until the tendon was able to be subluxed. With gentle traction, it was released from the glenoid by following the tendon posteriorly with a pair of curved dissecting scissors and cut at the level of the bone (Figure 1C).

The tendon was longitudinally split along its longitudinal fibers with a #15 blade with care taken to ensure a partial-thickness cut (Figure 1E-F). The tendon was then inverted inside-out (inverse tubularization) using a 6-0 prolene suture on a taper needle in two suture steps: First, an inverting simple interrupted stitch maintained inversion of the tendon at the musculotendinous junction. Second, a modified Kessler stitch was placed at the proximal tendon end, in an inverted configuration, and tied. This Kessler stitch was then used to pass the tendon through a bone tunnel drilled using a 20 G needle in an anterior-posterior fashion in the proximal humerus, lateral to the bicipital groove (Supplemental Video 1). A microvascular clip was placed on the tendon to complete a suspensory fixation in a length-controlled fashion by gently pulling the biceps muscle belly with the arm in 90 degrees of flexion. This was done while avoiding musculotendinous junction tissue involvement within the tunnel. Extra tendon and suture were removed. Fascia was closed in layers using 3-0 resorbable Vicryl suture. All animals received one dose of extended-release buprenorphine and permitted ad lib cage activity.

### Biomechanical testing

All specimens were freshly collected at the appropriate timepoint and underwent a single freeze-thaw cycle in −80°C storage to standardize biomechanical testing. Thawed specimens were cleaned of extraneous muscle soft tissue until only the tenodesed biceps tendon and intact humerus remained. The suspensory vascular clip was also removed from the opposite cortex to avoid interfere with strength testing of the repair. The humeri were potted in Bondo, and the tendon was mounted in a 320-grit sandpaper sandwich with cyanoacrylate glue as previously published^7^. Care was taken to keep soft tissues intact. The specimens were mounted in an Instron 8872 testing with a 50 N load cell for pull-to-failure testing using a displacement-control method at 2 mm per minute. All specimens were preloaded to 1 N prior to initiation of pull-to-failure. Peak failure forces were recorded and averaged. Failure mode was also recorded.

### Histology

Rat humeri were decalcified with EDTA. Tissues were fixed in zinc for 48 hours at room temperature and washed with PBS. Blocks were embedded in paraffin sagittally and sectioned at a thickness of 7 μm. Slides were subsequently deparaffinized with a xylene and ethylene glycol monomethyl ether (EGME) ladder before tinctorial and immunohistochemical staining. For hematoxylin and eosin (H&E) staining, slides were incubated with hematoxylin for 10 minutes at room temperature, washed with distilled water, incubated with eosin for 3 minutes at room temperature.

To detect lubricin, antigen retrieval was performed for 5 minutes at room temperature. Slides were then blocked with 2.5% normal horse serum for 20 minutes at room temperature before incubated with a primary antibody against lubricin or mouse IgG1 overnight at 4C. A kit was used to detect mouse IgG according to manufacturer’s instructions, and alkaline phosphatase was developed with Warp Red Chromogen. Slides were counterstained with 1% toluidine blue for 3 minutes at room temperature, dried, dehydrated with a xylene ladder only, and cover-slipped with acrylic media.

For Picrosirius red and Alcian blue staining, slides were incubated with hematoxylin, rinsed with distilled water, stained with Alcian blue for 30 minutes at room temperature, stained with Picrosirius red for 1 hour at room temperature, and rinsed in acid water. All slides were dehydrated with EGME and xylene and coverslipped with acrylic media unless otherwise noted. Slides were imaged using a Zeiss Axioscan 7 Slide Scanner at 20X with visible light. For Picrosirius red slides, circular polarized light (cPOL) was additionally used to detect collagen fibril alignment.

### The Bone Tunnel Interface Healing Score (IHS)

A new histological scoring system was created to semi-quantitatively measure and compare bone tunnel healing between surgical groups. The IHS was modified based on previously described rubrics^9,10^. Slides stained with H&E were graded on graft degeneration, collateral connection, and percent fibrous tissue using three fields of view. These readouts were chosen from existing literature because other categories were not applicable in our sections. Grading criteria ranged from 0 to 4 with integers as possible scores, and a 3×3 grid showing representative scores was given to blinded scorers (Table 1). Three 100 μm x 100 μm representative images from the healing interface for each specimen were presented to three blinded scorers who then graded each image independently. Only specimens that had the bone tunnel in-plane were included for analysis. The intraclass correlation coefficient (ICC) was determined to be 0.80 for all scores.

**Table 1.** Tendon-Bone Interface Healing Score (IHS) Rubric. Scoring rubric with criteria (left) and 3×3 grid showing representative scores of graft degeneration, collateral connection, and fibrous tissue (right).

|  | Grade 0 | Grade 2 | Grade 4 |
| --- | --- | --- | --- |
| **Graft degeneration** | Severe degeneration with <25% graft remaining and abundant empty space | Moderate degeneration with <50% of graft remaining | No graft degeneration |
| **Collateral connection** | 0% graft-to-bone healing interface | <50% graft-to-bone healing interface | 100% direct graft-to-bone connection with clear Sharpey fiber formation |
| **Fibrous tissue** | >75% fibrous tissue between graft remnant and cortical bone front | >50% and <75% fibrous tissue between graft remnant and cortical bone front | No fibrous tissue with 100% direct graft-to-bone connection |

### Immunohistochemical analyses

Images were imported and changed into 8-bit color in ImageJ. The distal bone-tendon interface was identified, and the region of interest (ROI) was defined as 100 μm proximal to the humeral head closer to the tenodesis aperture. This distal interface was chosen as the ROI, as it is the compression side of the tunnel. 100 μm was chosen to include the healing interface while excluding the endotenon. Percent area and signal intensity of the ROI were recorded. The same threshold level was used to benchmark images based on a negative control. Only specimens that had the bone tunnel in-plane were included for analysis.

### Statistical analyses

Data are reported as mean ± standard deviation and analyzed using unpaired t-tests. Significance was determined at p<0.05. Sample size was chosen based on power analyses from previous studies^7^. All statistics were performed using GraphPad Prism software or Stata.

## RESULTS

### Tendon inversion increases maximum tensile strength and stiffness

After mounting the tissue in our device (Figure 2A), peak failure force was measured. Modes of failure were also recorded (Table 2). During dissection, we noted two tendon inversion specimens had early tendon failure resulting in a Popeye deformity. One of the control specimens had uncharacteristic adhesion formation to the pectoralis insertion. These samples were excluded from our analysis. While there was no difference in first peak failure between the groups, there was a 27% increase in maximum peak failure in the tendon inversion group (11.0 ± 2.9 N) compared to the control group (8.6 ± 1.8 N) (Figure 2B). This suggests that tendon inversion has the capacity to increase tendon fixation strength.

**Table 2.** Biomechanical Testing Modes of Failure. Table summarizing biomechanical data with modes of failure. Rupture defined as either surface or midsubstance failure.

|  | First peak failure | First high peak | Max peak | Rupture | Tunnel pullout |
| --- | --- | --- | --- | --- | --- |
| **Control** | 3.91 | 6.34 | 8.61 | 11 | 1 |
| **Inversion** | 4.57 | 6.65 | 10.97 | 7 | 5 |

**Figure 2.**
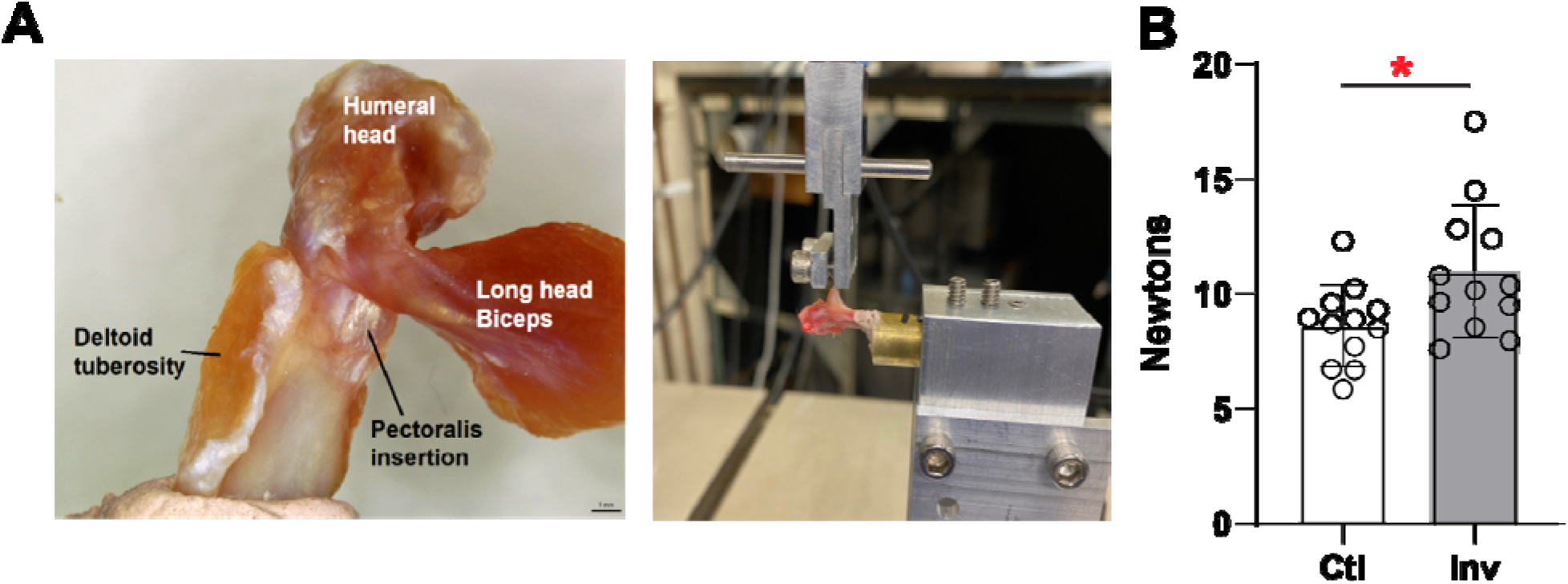
Tendon inversion increases maximum tensile strength and stiffness. (A) Surgical specimen (left) into the apparatus used to measure tendon failure force and stiffness (right). (B) Peak tendon failure force from control and inversion surgery. *p<0.05 by unpaired t-test.

### Tendon inversion decreases lubricin expression at the bone-tendon interface and enhances bone tunnel healing

Routine histology revealed direct intratendinous fascicular healing against the bone tunnel walls with evidence of mechanically-oriented fibrils with insertion into bone resembling Sharpey fibers (Figure 3A).

**Figure 3.**
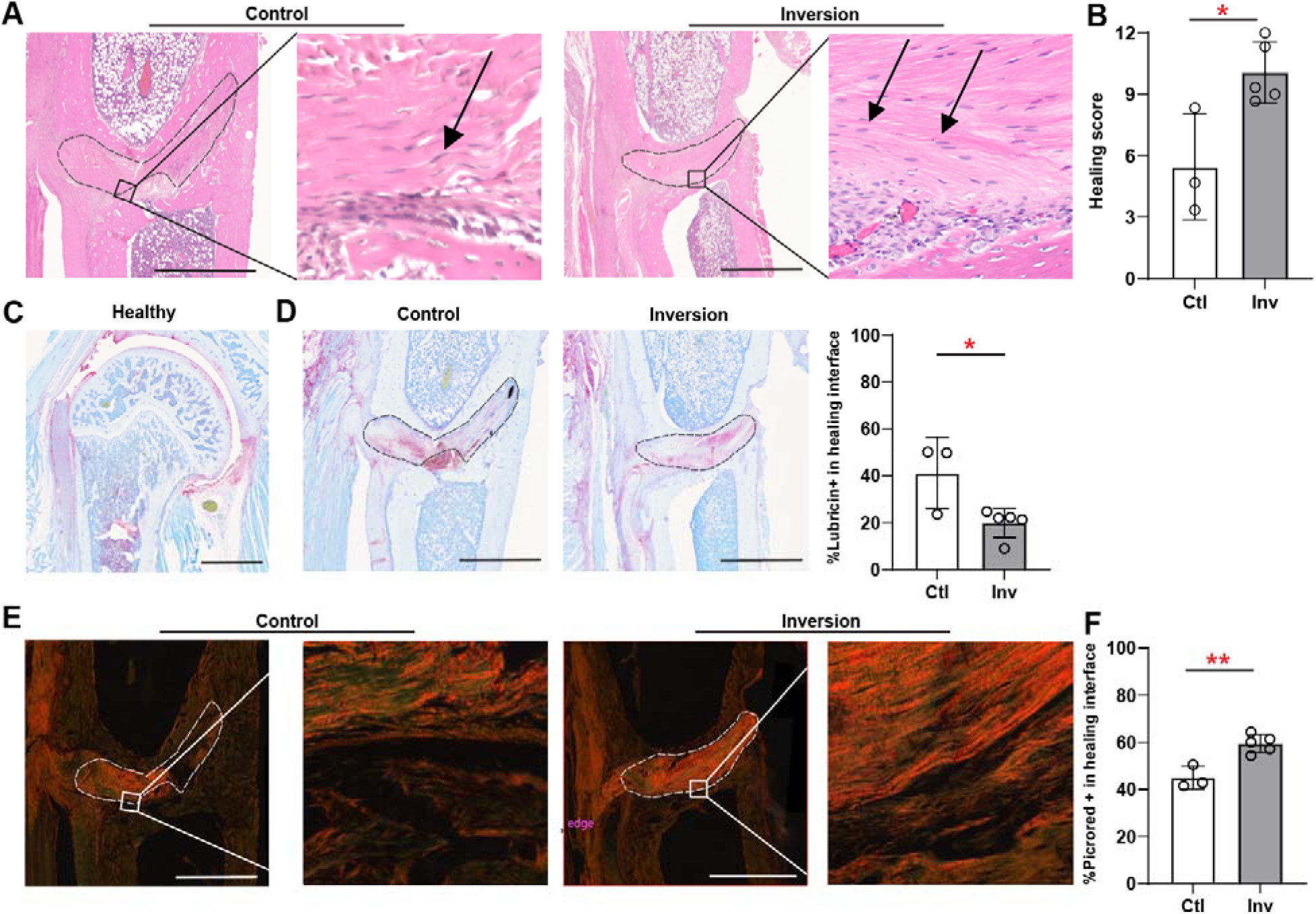
Tendon inversion decreases lubricin expression at the bone-tendon interface and enhances bone tunnel healing. (A) Hematoxylin and eosin (H&E) stained sections of control (left) and inversion surgery (middle) at 20X with additional magnification underscoring Sharpey fiber formation (right, inversion) or lack thereof (left, control). (B) Bone tunnel healing score quantification of control and inversion surgery. (C) Lubricin on a healthy humeral head and shaft detected by Warp Red Chromogen counterstained with Toluidine blue. (D) Lubricin on control (left) and inversion surgery (middle) at 20X on brightfield imaging with quantification of percent-positive lubricin in the bone tunnel (right). (E) Picrosirius red on control and inversion surgery at 20X under circular polarized light with additional magnification. (F) Quantification of percent-positive Picrosirius red at the bone-tendon healing interface. *p<0.05 and **p<0.01 by unpaired (B, D, F) t-test. Scale bars measure 2 mm (A, C, E). Dashed lines represent the tendon graft.

On histologic scoring of bone tunnel healing, animals that underwent control surgery showed moderate IHS scores (5.4 ± 2.6) whereas animals that had tendon inversion showed near two-fold increased IHS scores based on graft degeneration, collateral connections, and fibrous tissue infiltration (10.1 ± 1.5) (Figure 3B).

Immunohistochemical studies revealed lubricin on cartilaginous surfaces as positive control as previously demonstrated^10^ (Figure 3C). After tendon inversion, lubricin expectedly localized to the endotenon and had decreased expression at the healing interface compared to control specimens (41.2% vs. 19.9%) (Figure 3D). These findings suggest that biomechanical and bone tunnel healing differences are associated with decreased lubricin expression at the healing interface.

Collagen fibril deposition and alignment were measured. Type III collagen is initially formed at the wound bed and gradually replaced by stronger, aligned type I collagen^15^. Control specimens had smaller percent-positive Picrosirius red area at the interface compared to inversion specimens (45.0% vs. 59.5%) (Figure 3E-F). This was independent of birefringence intensity. Together, our data suggest that decreased lubricin expression at the bone tunnel interface may, in part, contribute to superior biomechanical and bone tunnel healing differences seen in our tendon inversion model.

## DISCUSSION

Here, we establish a new model of rat biceps tenodesis whereby the tendon construct is inverted, exposing intratendinous fascicles and limiting epitendinous tissue at the bone interface. We find that this procedure leads to increased maximum peak failure force of the healed repair. We also show that tendon inversion enhances histologic bone tunnel healing and collagen fibril alignment while decreasing lubricin expression at the interface. Our results suggest that purposeful native tendon alterations may improve tendon-to-bone healing.

Most research to date on tendon-to-bone repair has focused on technique and implant design to reduce the risk of surgical failure. While these efforts are important to streamline procedures, the biological mechanisms behind tissue healing following injury and surgery are not as well understood. As such, dissecting these healing processes can better inform therapeutic strategies. Tendon attaches to bone via an enthesis, a unique fibrocartilaginous structure that is affected by a variety of factors including intrinsic biology and mechanical load^13^. Following injury, the tendon must be reattached to the bone at the enthesis for proper healing and subsequent increased function. Indeed, biomarkers such as growth factors, cytokines, and other small molecules are dysregulated in acute and chronic tendon disease including interleukin (IL)-1β, cyclooxygenase (COX)-2, tumor necrosis factor (TNF)-α, and matrix metalloproteinase (MMP) subgroups^14^. Fibrovascular scar tissue develops, and coupled with inflammatory and remodeling factors, this combination complicates the already-challenging task of fixing the compliant tendon to stiff bone^13^. As such, recapitulating this natural biomechanical gradient may improve longevity of the tissue repair and minimize subsequent failure. Recent work has highlighted individual pathways involved in tissue repair, such as hedgehog activation, targeted nuclear factor κβ (NF-κβ) inhibition, and fibroblast growth factor (FGF) augmentation for improved tendon-to-bone healing^14-16^. Studies interrogating bioactive scaffolds and their properties have also demonstrated promise in bolstering tendon-to-bone repair^17^.

In addition to the environmental milieu, the biology of tendons and their native sliding function may interfere with bone tunnel healing. Lubricin is one such glycoprotein involved in lubricating tendons. Taguchi and colleagues previously reported that exogenous lubricin enhances tendon gilding ability in a canine flexor digitorum profundus model^20^. Further, models of lubricin knockout mice demonstrated increased gliding resistance, underscoring lubricin’s importance in intrasynovial tendon lubrication^21^. As such, it is plausible that disrupting lubricin at the repair interface may improve tendon to bone healing. Surgical dogma stipulates scuffing smooth tendon surfaces to improve tendon-to-bone healing or instead performing bone surface tenodesis without a tunnel, thereby permitting the tendon to scar against periosteum. Both manipulations achieve similar results by exposing the endotenon and fibrils directly to the healing interface^22^. Modern methods typically use hamstring allografts with intact epitenon; there is little consideration for the tissue quality or type. Other techniques utilize tunnel fixation with interference screws; suspensory fixation, which creates bone sockets for graft passage; and surface fixation^23-24^. On our review of the literature, however, tendon inversion has not been previously described.

Our study does have limitations. While our data shows significant improvement in our rat model, these findings may not directly translate to human tendons. Rats were chosen as the model due to anatomic similarities to humans. However, quadruped species have different force distributions compared to human biceps. There is therefore a need to test for injurious effects of tendon inversion on human tendon tensile strength at time zero before introducing this technique to patients. Further, while we have shown an association between lubricin expression and functional and healing readouts, our data do not speak to causality. Future studies should address the importance of lubricin, and other glycoproteins, with in vivo lubricin blockade or genetic models.

Together, our findings demonstrate that a novel surgical method of tendon inversion improves tendon-to-bone repair healing, which is associated with decreased lubricin expression at the tendon-bone interface. Future efforts should investigate safe human application of lessons learned from this work, as well as lubricin manipulation in tendon-to-bone healing.

## FIGURE LEGENDS

**Supplemental Video 1. Surgical Approach to the rat biceps tendon**.

## ACKNOWLEDGEMENTS

**Acknowledgements:** The authors thank the Orthopaedic Surgery Resident Research Committee at the Icahn School of Medicine at Mount Sinai for helpful discussion.

## Funding

This work was funded by the American Foundation for Surgery of the Hand – Basic Science Grant and a Mount Sinai Orthopaedic Resident Research Award.

## AUTHOR CONTRIBUTIONS

Conceptualization: GTC, PJC, MRH

Performed experiments and acquired data: GTC, TML, DCB, VA, PN, DL, HRF, AJH

Data analyses and interpretation: GTC, TML

Methodology and reagents: PN, DL, DML

Supervision: AHH, PJC, LMG, MRH

Writing–original draft: GTC, TML Writing–reviewing and editing: All authors

## Notes

### Competing Interest Statement

The authors have declared no competing interest.

